# Rovaca: A highly optimized variant caller for rapid and robust germline DNA analysis

**DOI:** 10.1101/2025.10.19.677660

**Authors:** Qiwen Zheng, Longhui Yin

## Abstract

The Genome Analysis Toolkit (GATK) HaplotypeCaller is the gold standard for germline variant detection, but its computational intensity presents a significant bottleneck in large-scale genomic analyses. While accelerated solutions exist, they often require specialized hardware or are proprietary, limiting accessibility.

We present Rovaca, a novel, pure software variant caller implemented in C++ that dramatically accelerates germline analysis on standard CPU hardware. By preserving the core mathematical models of GATK HaplotypeCaller, Rovaca achieves nearly identical output, ensuring high concordance and facilitating adoption. Performance gains are achieved through a sophisticated producer-consumer multi-threading architecture and targeted micro-architectural optimizations of the PairHMM algorithm using AVX-512 instructions. Benchmarks on standard GIAB datasets show Rovaca is 57-76× faster for Whole-Genome Sequencing and 30× faster for Whole-Exome Sequencing compared to GATK. The F1-scores for both SNPs and INDELs consistently differ by less than 0.01%, demonstrating exceptional accuracy. Rovaca is a deterministic, robust, and cost-efficient tool that removes the computational barrier for high-throughput germline variant analysis.

Rovaca is open-source and available at https://github.com/ZephyRoy/Rovaca.

## 1 Introduction

The advent of next-generation sequencing (NGS) technologies has revolutionized the field of human genetics, enabling comprehensive analysis of the entire genome (Whole-Genome Sequencing, WGS) or its protein-coding regions (Whole-Exome Sequencing, WES) [6, 13]. Due to rapidly declining costs and increasing throughput, WGS and WES are no longer confined to research laboratories; they have become integral tools in clinical diagnostics for identifying the genetic basis of Mendelian disorders, characterizing rare or novel diseases, and informing personalized medicine [4, 1, 18, 10].

However, the massive volume of data generated by NGS platforms presents a significant computational challenge. A standard workflow involves aligning sequencing reads to a reference genome, preprocessing the alignment file, and finally performing variant calling—the identification of single nucleotide polymorphisms (SNPs) and short insertions/deletions (indels) [11]. This entire process is computationally intensive, often requiring tens of hours to analyze a single genome, which is a critical bottleneck in time-sensitive clinical scenarios and large-scale population studies [16].

For years, the Genome Analysis Toolkit (GATK), developed by the Broad Institute, has been the gold standard for germline variant calling, with its HaplotypeCaller algorithm recognized for high accuracy and sensitivity [17, 3]. The GATK Best Practices workflow is the benchmark against which other tools are measured. However, GATK’s primary limitation has been its computational performance. This performance gap has spurred the development of numerous alternative solutions (Table1).

**Table 1:** Comparison of modern germline variant calling tools.

| Tool | Algorithm Basis | Hardware | License | Key Feature |
| --- | --- | --- | --- | --- |
| GATK HC | Haplotype Assembly | CPU | Open Source | Gold standard |
| DeepVariant | Deep Learning | GPU | Open Source | High accuracy |
| DRAGEN | Hardware-specific | FPGA | Commercial | Ultra-fast |
| Strelka2 | Custom statistical | CPU | Open Source | Fast somatic/<br>germline |
| Sentieon | GATK re-implementation | CPU | Commercial | GATK-identical |
| **Rovaca** | **GATK re-implementation** | **CPU** | **Open Source** | **GATK-identical, 50x faster** |

These alternatives can be broadly categorized into hardware-accelerated and software-based solutions. Hardware-based approaches like Google’s DeepVariant [12] and Illumina’s DRAGEN platform achieve dramatic speedups but require specialized, often costly, hardware (GPUs/FPGAs). In parallel, software-optimized tools like Strelka2 [9] and Sentieon DNASeq [8] have re-implemented core algorithms in efficient languages like C/C++. However, they come with trade-offs: Strelka2’s output differs from GATK, creating a barrier for adoption, while Sentieon is a commercial product requiring a license.

Here, we introduce Rovaca, a new, fast, and robust variant caller that overcomes these limitations. Rovaca is a pure software solution, implemented in C++, that strictly adheres to the mathematical methods of the GATK HaplotypeCaller. By employing a specifically designed multi-threading architecture and novel AVX instruction optimizations, Rovaca delivers a 50- to 70-fold speed improvement over GATK HC on standard CPUs. It maintains extremely high concordance with GATK’s results, requires no specialized hardware, and is open-source. This paper describes the architecture of Rovaca and presents a comprehensive benchmark, demonstrating its performance as a fast, accurate, and cost-efficient solution for today’s genomic analysis needs.

## 2 Results

To evaluate the performance of Rovaca, we conducted a series of comprehensive benchmarks focusing on two key areas: computational performance (speed and scalability) and variant calling accuracy (consistency with the established standard). We compared Rovaca against the GATK HaplotypeCaller (HC) using publicly available benchmark datasets from the Genome in a Bottle (GIAB) consortium.

### Computational Performance and Scalability

A primary design goal of Rovaca is to offer a substantial reduction in the time required for variant calling on standard CPU-based hardware. We first assessed its multi-threading performance and then compared its runtime against GATK HC on both Whole-Genome Sequencing (WGS) and Whole-Exome Sequencing (WES) data.

### Multi-threading Scalability

We first assessed the scalability of Rovaca by running it on the HG001 35X WGS dataset with an increasing number of CPU threads. As illustrated in Figure1, Rovaca’s runtime decreases substantially as the number of threads increases, closely following the ideal inverse relationship. The most significant performance gains are observed when scaling up to approximately 50 threads. Beyond this point, while the runtime continues to decrease, the speedup becomes less pronounced as it approaches a performance plateau. Conversely, peak memory usage demonstrates a near-linear increase with the number of allocated threads. This analysis indicates that Rovaca is highly efficient at parallelizing the variant calling workload and suggests that users can select an optimal thread count to balance speed with available computational resources.

**Figure 1.**
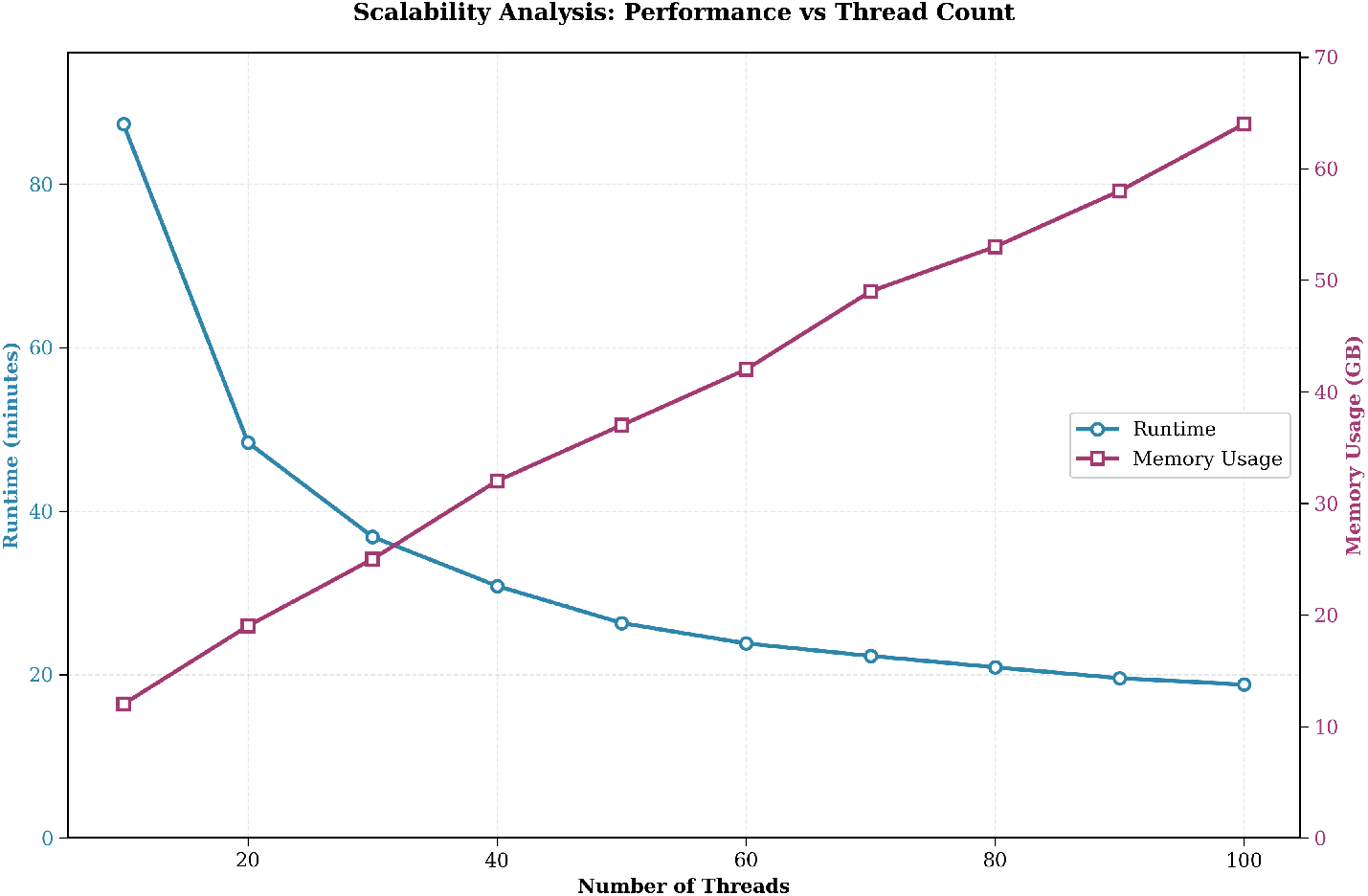
Time and memory consumption for Rovaca on the HG001 35X WGS dataset across a range of thread counts. The blue line indicates runtime in minutes (left y-axis), and the dark pink line indicates peak memory usage in gigabytes (right y-axis).

### Runtime Comparison on WGS and WES Data

To benchmark Rovaca against the current standard, we compared its runtime to GATK HaplotypeCaller on seven different WGS samples at approximately 35X coverage. Rovaca demonstrated a dramatic speedup across all samples, ranging from **57x to 76x** (Figure2). For instance, on the HG004 sample, Rovaca completed the analysis in just 18.2 minutes, whereas GATK HC required 1385.7 minutes (a 76-fold improvement). On average, GATK HC took approximately 22 hours to process a sample, while Rovaca completed the same task in about 20 minutes.

**Figure 2.**
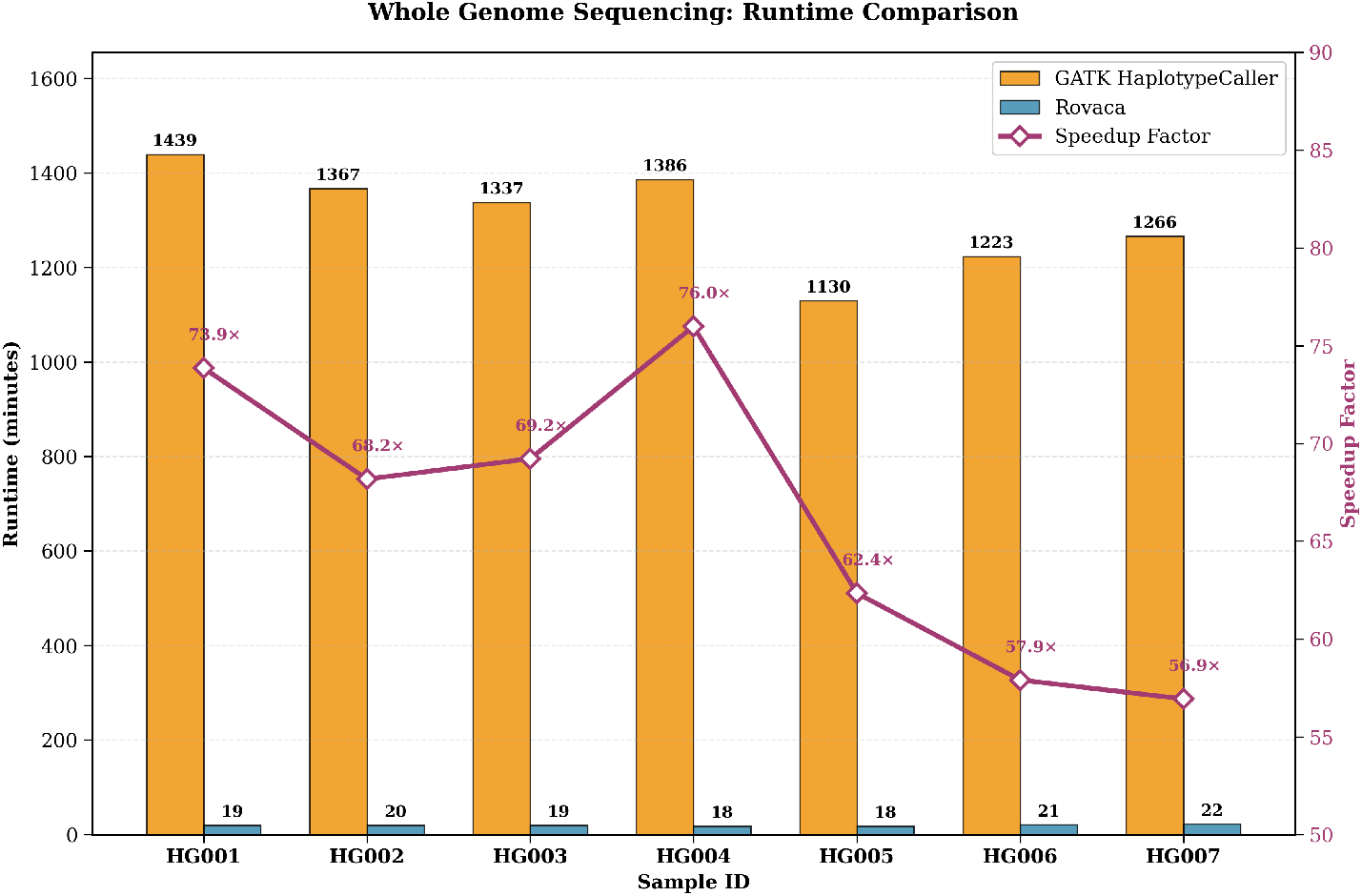
Runtime comparison for Rovaca and GATK HC on seven 35X WGS samples with VCF output. Orange bars represent GATK HC runtime in minutes, blue bars represent Rovaca runtime in minutes (left y-axis), and the dark pink line indicates the speedup factor (right y-axis).

Similar performance gains were observed for WES data. On the same seven samples, Rovaca achieved an average speedup of approximately **30-fold** (Figure3). It consistently processed each WES sample in about 1.3 minutes, compared to GATK HC’s average runtime of approximately 40 minutes per sample. These results confirm Rovaca’s efficiency in significantly reducing the computational time for both WGS and WES analyses.

**Figure 3.**
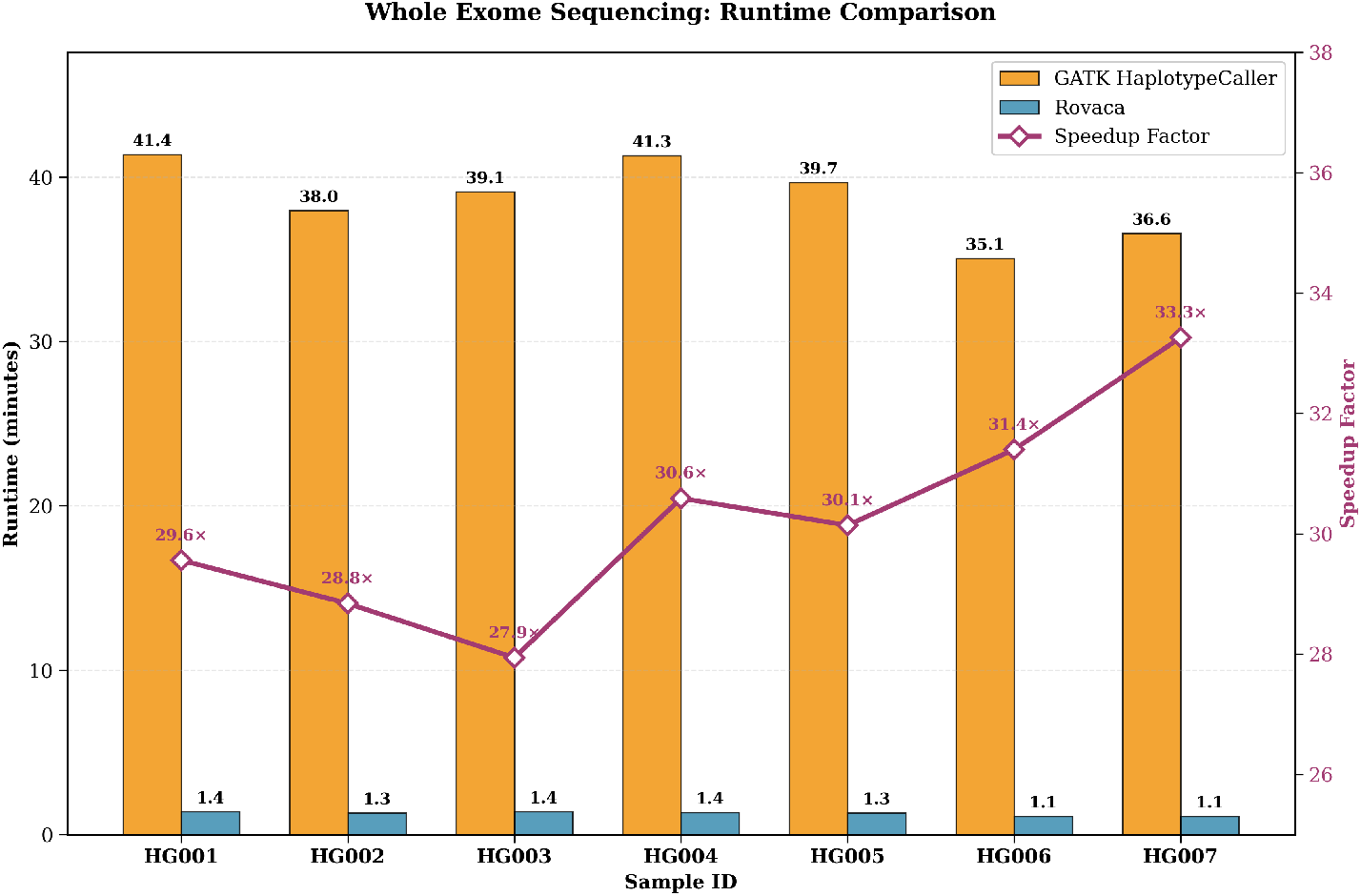
Runtime comparison for Rovaca and GATK HC on seven WES samples with VCF output. Orange bars represent GATK HC runtime, blue bars represent Rovaca runtime, and the dark pink line indicates the speedup factor.

### Accuracy and Consistency Analysis

While speed is a primary advantage, it is crucial that it does not come at the expense of accuracy. We used the hap.py tool to perform a rigorous comparison of the variant calls from Rovaca and GATK HC against the GIAB truth sets. Our analysis demonstrates that Rovaca produces a variant call set that is highly concordant with GATK HC.

As shown in Table 2, across multiple samples (HG001, HG002, HG003, HG004), reference genomes (GRCh37, GRCh38), and sequencing strategies (WGS, WES), the F1-scores for both SNPs and INDELs were consistently within 0.01

**Table 2:**
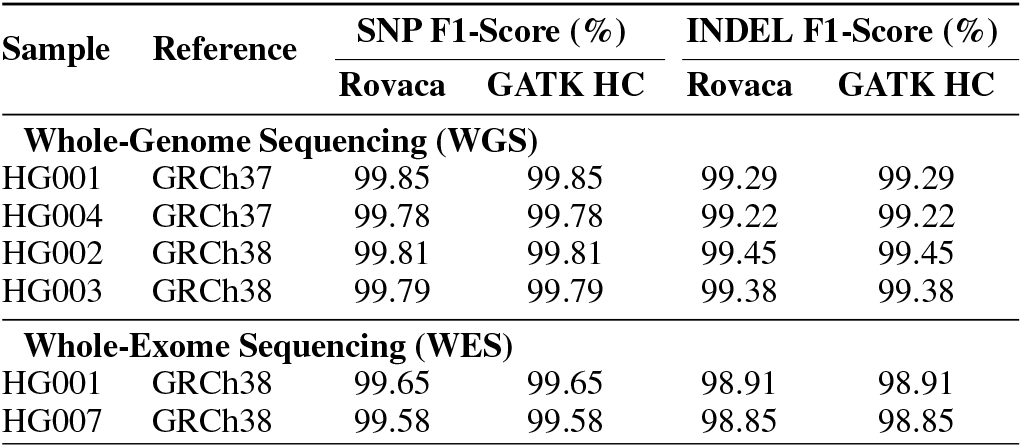
Accuracy and Concordance Analysis. F1-scores for Rovaca and GATK HC were compared against GIAB truth sets for various samples, reference genomes, and sequencing types. The difference between the F1-scores is negligible.

| Sample | Reference | SNP F1-Score (%) |  | INDEL F1-Score (%) |  |
| --- | --- | --- | --- | --- | --- |
|  |  | Rovaca | GATK HC | Rovaca | GATK HC |
| Whole-Genome Sequencing (WGS) |  |  |  |  |  |
| HG001 | GRCh37 | 99.85 | 99.85 | 99.29 | 99.29 |
| HG004 | GRCh37 | 99.78 | 99.78 | 99.22 | 99.22 |
| HG002 | GRCh38 | 99.81 | 99.81 | 99.45 | 99.45 |
| HG003 | GRCh38 | 99.79 | 99.79 | 99.38 | 99.38 |
| Whole-Exome Sequencing (WES) |  |  |  |  |  |
| HG001 | GRCh38 | 99.65 | 99.65 | 98.91 | 98.91 |
| HG007 | GRCh38 | 99.58 | 99.58 | 98.85 | 98.85 |

These results establish that Rovaca is not only significantly faster than GATK HaplotypeCaller but also produces a highly concordant set of variant calls, making it an accurate and efficient alternative for germline variant discovery.

## 3 Discussion

Rovaca is a highly optimized, pure software solution designed to address the critical need for rapid and accurate germline variant calling from NGS data. A key aspect of its design is its versatility and adherence to community standards, making it suitable for a wide range of genomic analysis pipelines. It seamlessly processes both Whole-Genome Sequencing (WGS) and Whole-Exome Sequencing (WES) data, with the latter facilitated by the standard inclusion of a BED file to define target regions. Furthermore, Rovaca supports the generation of output in both standard VCF and genomic VCF (GVCF) formats. This dual-format capability satisfies the requirements for both single-sample analysis and large-scale cohort studies, which rely on joint genotyping from GVCFs. These features ensure that Rovaca can be easily integrated into most existing workflows, serving the needs of a broad user base.

The most striking contribution of Rovaca is its dramatic improvement in computational speed while running on standard x86-CPU-based systems. Our benchmarks demonstrate a remarkable 50- to 70-fold speedup for WGS and an approximately 30-fold speedup for WES compared to the widely-used GATK HaplotypeCaller. This acceleration is a direct result of Rovaca’s ground-up C++ implementation, which features a sophisticated multi-threading architecture based on a producer-consumer model, and targeted algorithmic optimizations. By utilizing shared thread and memory pools, Rovaca minimizes I/O bottlenecks and maximizes CPU utilization. As our scalability analysis shows, Rovaca effectively utilizes available CPU cores, with performance gains increasing steadily with the number of threads. However, as the number of threads exceeds 50, the performance gains begin to plateau while memory consumption continues to rise. This observation provides a practical recommendation for users: they should select a thread count based on their specific hardware environment to optimize the trade-off between speed and resource allocation.

Crucially, this substantial acceleration is achieved without compromising the accuracy and reliability of variant detection. Our extensive comparisons using the GIAB benchmark datasets show that Rovaca’s variant calls exhibit extremely high concordance with those produced by GATK HC. Across multiple samples, reference genomes (GRCh37 and GRCh38), and sequencing strategies (WGS and WES), the F1-scores for both SNPs and INDELs were consistently within 0.01% of those from GATK HC. This is a direct result of our deliberate choice to adhere strictly to the underlying mathematical models of the GATK HaplotypeCaller, ensuring that the biological integrity of the results is preserved. Users can therefore benefit from the significant speedup without questioning the validity of the output.

Another critical feature of Rovaca is its deterministic nature. Unlike some parallel computing frameworks that can introduce stochastic variability, Rovaca produces identical results on identical inputs across multiple runs. This run-to-run consistency is paramount in clinical and regulated environments where reproducibility is a strict requirement for validation and diagnostics.

The combination of speed, accuracy, and accessibility makes Rovaca a powerful tool for a wide range of applications. In large-scale population genomics projects, the ability to process thousands of genomes rapidly and cost-effectively is essential. In the clinical setting, particularly for the diagnosis of rare diseases in newborns or in time-sensitive oncology cases, reducing the analysis time from days to hours, or hours to minutes, can be transformative. By operating on standard x86-CPU architecture, Rovaca eliminates the need for specialized and expensive hardware like GPUs or FPGAs required by other accelerated solutions such as DeepVariant or DRAGEN, thereby democratizing high-throughput genomic analysis for a broader range of research and clinical laboratories.

While Rovaca represents a significant advance, we acknowledge its current limitations and see clear avenues for future development. The current version is optimized specifically for germline short-variant (SNP and INDEL) detection. Future work will focus on expanding its capabilities to include somatic variant calling for cancer genomics, which presents a different set of challenges, as well as the detection of larger structural variants (SVs) and copy number variations (CNVs) by combining both short read sequencing and long read sequencing.

In conclusion, Rovaca successfully addresses the critical need for a fast, cost-efficient, and accurate variant caller for human germline DNA analysis. By combining a highly parallelized software architecture with targeted algorithmic optimizations, it provides a dramatic speedup over the GATK HaplotypeCaller while maintaining exceptional concordance. Its reliance on standard CPU hardware makes it an accessible and scalable solution for both research and clinical settings. Rovaca represents a significant step forward in democratizing high-throughput genomic analysis, enabling researchers and clinicians to move from raw sequencing data to meaningful biological insights more rapidly than ever before.

## 4 Methods

### 4.1 Datasets

We utilized seven publicly available benchmark datasets from the Genome in a Bottle (GIAB) consortium (HG001-HG007) for both Whole-Genome Sequencing (WGS) and Whole-Exome Sequencing (WES) evaluations[2]. The raw paired-end FASTQ files were obtained from the National Center for Biotechnology Information (NCBI) repository[5]. For the final accuracy benchmark, the corresponding high-confidence truth set VCF files and BED files were also obtained from the GIAB project hosted on the NCBI website. All accuracy assessments were performed using the hap.py tool from Illumina[7].

### 4.2 Hardware

All computational benchmarks were executed on a machine equipped with an Intel(R) Xeon(R) Gold 6248 CPU @ 2.50GHz and 384GB of memory.

### 4.3 Variant Calling Workflow

Our analysis pipeline was designed to adhere strictly to the established GATK best practices workflow[15, 14]. Before variant calling, this process involves three main preprocessing stages: (1) alignment of raw FASTQ reads to the reference genome using BWA-MEM, (2) marking of duplicate reads using GATK MarkDuplicates, and (3) Base Quality Score Recalibration (BQSR) to generate an analysis-ready BAM file.

The only modification to this standard pipeline was the replacement of the GATK HaplotypeCaller (version 4.6.0.0) with Rovaca for the final variant calling step. This ensures that our performance and accuracy comparisons are conducted under equivalent conditions, isolating the impact of the variant caller itself.

### 4.4 Rovaca Software Architecture and Optimizations

Rovaca is engineered from the ground up as a pure software solution implemented in C++, designed for maximum performance on commodity x86-CPU-based systems. Its architecture is fundamentally based on a highly efficient producer-consumer model, which utilizes a shared thread and memory pool to enable dynamic resource allocation and minimize computational overhead. The overall software architecture is depicted in Figure4.

**Figure 4.**
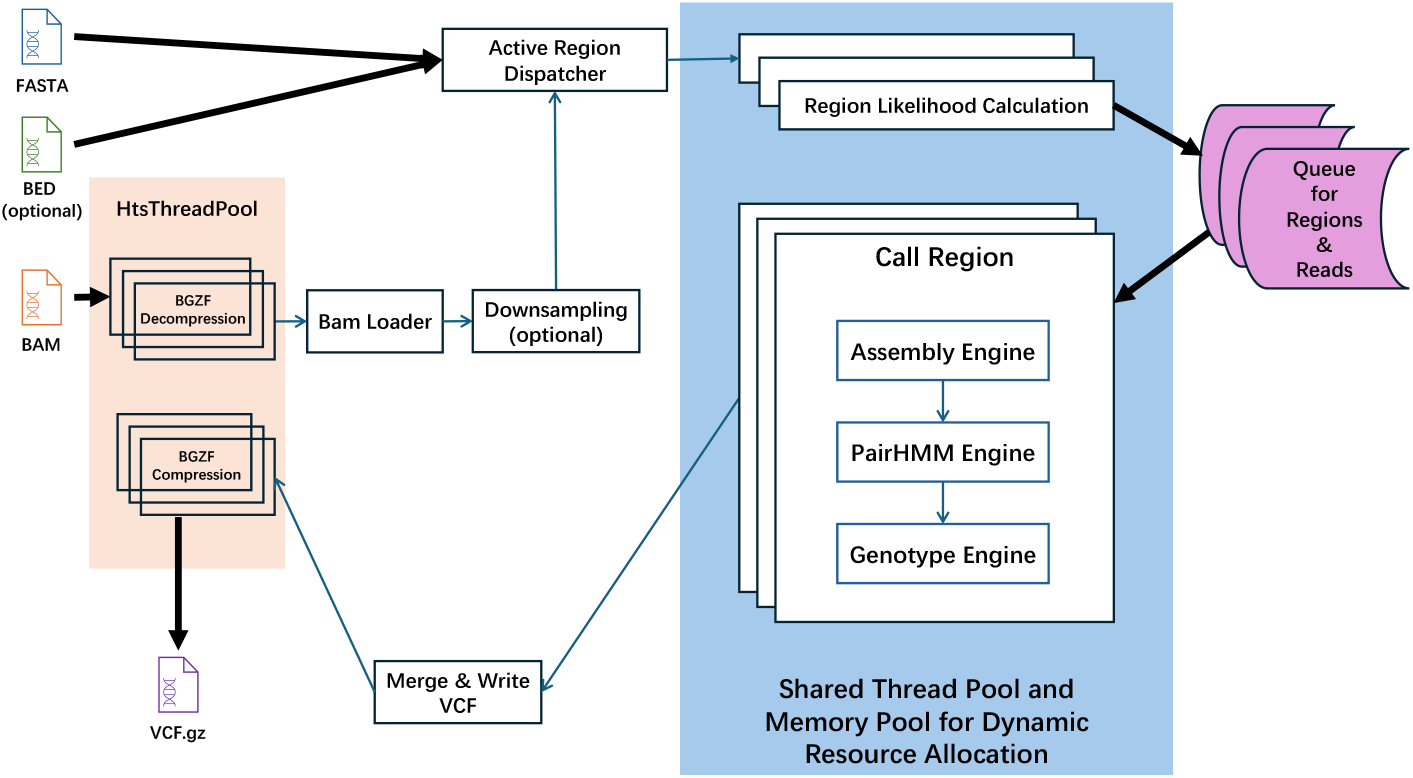
The overall software architecture design of Rovaca, illustrating the producer-consumer pattern and the flow of data through different modules. This design facilitates dynamic resource allocation from a shared thread and memory pool.

The workflow initiates with the parallelized processing of the input alignment data from a BAM file. An HtsThreadPool manages the BGZF decompression, feeding reads into a Bam Loader. This loader extracts the necessary read information, which is then passed to the Active Region Dispatcher. This dispatcher module, using the reference FASTA and an optional BED file to define the analysis scope, identifies genomic regions with sufficient evidence of potential variation. For regions with excessive coverage, an optional downsampling step can be applied to normalize read depth before dispatching.

Once an active region is identified, its genomic coordinates and all associated reads are bundled into a computational task. This task is then submitted to a central Queue for Regions & Reads, which serves as the central buffer between the producer and consumer stages. The core computational modules, including Region Likelihood Calculation and the multi-stage Call Region engine, act as consumers. These modules draw tasks from the queue and process them in parallel. The Call Region engine itself is composed of three key components: the Assembly Engine for local de novo assembly of haplotypes, the highly-optimized PairHMM Engine for read-haplotype likelihood calculation, and the Genotype Engine for final variant determination.

A central innovation in Rovaca’s design is the use of a single, shared thread pool and memory pool for all computationally intensive stages. This architecture facilitates dynamic resource allocation, where threads are not statically assigned but are drawn from the shared pool as tasks become available. When a task is completed, its resources are immediately returned to the pool, ensuring maximum CPU utilization and minimizing idle time. This design is the primary driver behind Rovaca’s high thread efficiency and comparatively low memory footprint.

To ensure proper genomic ordering in the final output, variant calls from completed regions are written to temporary, chromosome-specific files. Upon processing all regions, a final `Merge & Write VCF` step efficiently combines these intermediate files into a single, sorted, and BGZF-compressed VCF file (VCF.gz). This entire architecture is designed to minimize I/O operations and maximize in-memory processing, contributing significantly to Rovaca’s overall speed.

The primary performance bottleneck in the GATK HaplotypeCaller is the PairHMM algorithm, which is essential for calculating read-haplotype likelihoods. While Intel has previously provided an AVX-optimized implementation, we identified further opportunities for significant acceleration. Rovaca incorporates two major improvements to the PairHMM algorithm.

1. **Intelligent Read Batching Strategy for SIMD-Optimized PairHMM Computation** The intelligent batching strategy represents a novel approach to optimizing pairwise Hidden Markov Model (PairHMM) computations for genomic sequence alignment by leveraging the architectural characteristics of modern SIMD (Single Instruction, Multiple Data) processors, specifically AVX-512. This strategy addresses the fundamental challenge of efficiently utilizing 512-bit vector units when processing variable-length genomic reads, which traditionally suffer from significant computational inefficiency due to irregular memory access patterns and suboptimal vectorization. The core of this strategy employs a three-phase adaptive grouping algorithm that maximizes SIMD utilization through length-aware read clustering, while minimizing the computational overhead associated with padding and masking operations inherent in variable-length sequence processing. **Phase 1: Histogram-Based Length Distribution Analysis**. The algorithm begins by constructing a length histogram of all input reads, creating a statistical profile that enables optimal batch formation. This preprocessing step has *O*(*n*) complexity where *n* is the number of reads, and provides crucial insights into the data distribution that guide subsequent grouping decisions. **Phase 2: Homogeneous Length Batching**. The strategy prioritizes the formation of homogeneous batches containing exactly 16 reads of identical length (matching the AVX-512 vector width for single-precision floating-point operations). This approach eliminates the need for length-based masking operations during SIMD computation, achieving optimal computational efficiency. For length *ℓ* with count *c ≥* 16, the algorithm forms [*c/*16] complete batches, maximizing throughput for the most common read lengths. **Phase 3: Heterogeneous Length Consolidation**. For remaining reads that cannot form complete homogeneous batches, the algorithm employs a sophisticated neighborhood-based grouping strategy. It examines reads within a configurable length window (*±*5 nucleotides) and intelligently combines them to form mixed-length batches of size 16. This phase uses a bidirectional search algorithm that preferentially selects shorter reads first to minimize padding overhead, while ensuring that the maximum length difference within any batch remains bounded. The strategy incorporates adaptive thresholds (`k_read_group_count = 8`) that determine when heterogeneous batching becomes beneficial. This prevents the algorithm from creating suboptimal batches when the computational overhead of mixed-length processing exceeds the benefits of vectorization. The batching strategy is also tightly coupled with a memory layout optimization that arranges read data in a structure-of-arrays (SoA) format, enabling efficient SIMD loads and minimizing cache misses. Each batch is processed with a single haplotype across all 16 reads simultaneously, maximizing data reuse and computational intensity.
2. **Micro-architectural Optimization for SIMD Data Movement Efficiency** The second critical optimization addresses a fundamental inefficiency in SIMD data movement operations within AVX-512 registers, specifically targeting the reduction of Cycles Per Instruction (CPI) through strategic data layout manipulation. This optimization exploits asymmetric instruction latencies inherent in the x86-64 AVX-512 instruction set architecture. Our micro-architectural analysis of the original Intel AVX-512 implementation revealed a systematic performance bottleneck in the vector shifting operations required by the PairHMM dynamic programming algorithm. The original implementation employed a multi-step extraction process that required:
  1. **High-order bit extraction**: `_mm512_extractf32×8_ps(x.d, 1)` to extract the upper 256 bits
  2. **Scalar extraction**: `_mm256_extract_epi32(xhigh.i, SHIFT_CONST2)` to extract the highest 32-bit element
  3. **Complex permutation operations**: Multiple `VEC_PERMUTE, VEC_AND, and VEC_OR` operations for data rearrangement

This approach necessitates accessing the **most significant bits** of the vector register, which requires expensive inter-lane data movement and complex bit manipulation operations. Specifically, the `_mm256_extract_epi32`(v1, 7) operation incurs significant latency due to:

- **Cross-lane data dependency**: AVX-512 operates on independent 256-bit lanes, requiring inter-lane communication
- **High-latency extraction**: Extracting elements from higher indices requires additional routing through the shuffle units
- **Pipeline stalls**: The complex extraction sequence creates dependency chains that limit instruction-level parallelism

Our investigation revealed a critical asymmetry in AVX-512 microarchitecture: accessing the **lowest-order elements** (indices 0-3) of a vector register is substantially more efficient than accessing higher-order elements due to:

1. **Register file organization**: Lower indices map directly to the least significant portions of the physical register
2. **Reduced routing complexity**: No cross-lane data movement required
3. **Single-cycle extraction**: `_mm512_cvtss_f32` can extract the lowest float in a single cycle
4. **Simplified alignment**:`_mm512_alignr_epi32` with immediate count 1 performs efficient element-wise rotation

To exploit this micro-architectural characteristic without compromising algorithmic correctness, our implementation employs a **strategic data reversal** during the memory-to-register loading phase. This transformation ensures that the algorithmically “last” element (which needs to be extracted during vector shifting) is positioned at the **lowest index** (most efficient to access) in the SIMD register.

Algorithm 1 presents the pseudocode for this optimization:

The data reversal enables the replacement of complex extraction sequences with highly efficient single-instruction operations. Algorithm 2 shows the optimized vector shifting procedure:

This optimization delivers measurable improvements across multiple performance metrics:

- **Instruction Count Reduction**: 6-8 instruction reduction per iteration (from *∼*12 to *∼*4 instructions)
- **Latency Improvement**: Reduction from 8-12 cycles to 2-3 cycles per shift operation

#### Algorithm 1 Strategic Data Layout Reversal for SIMD Optimization

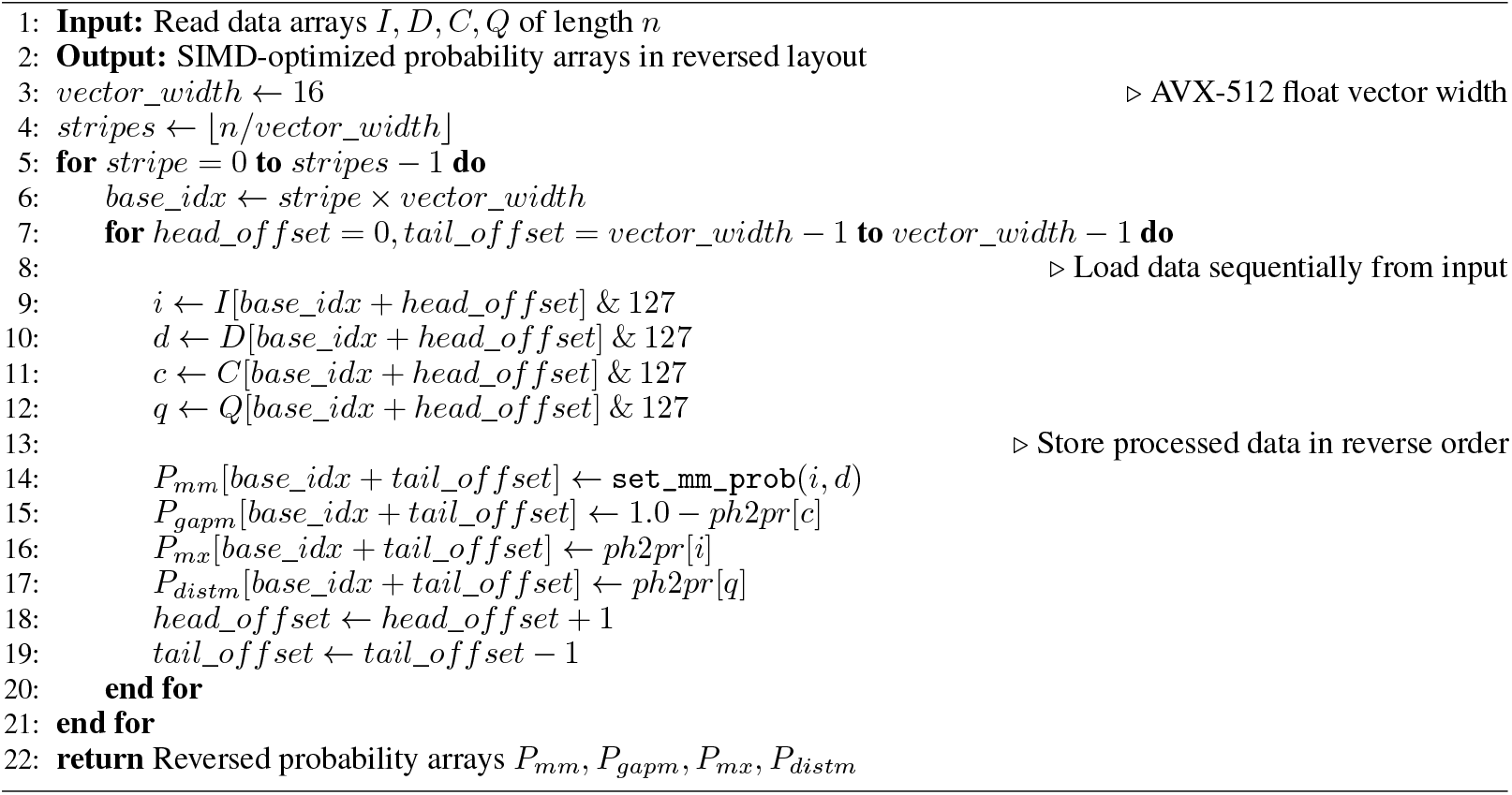

#### Algorithm 2 Optimized Vector Shift Operation

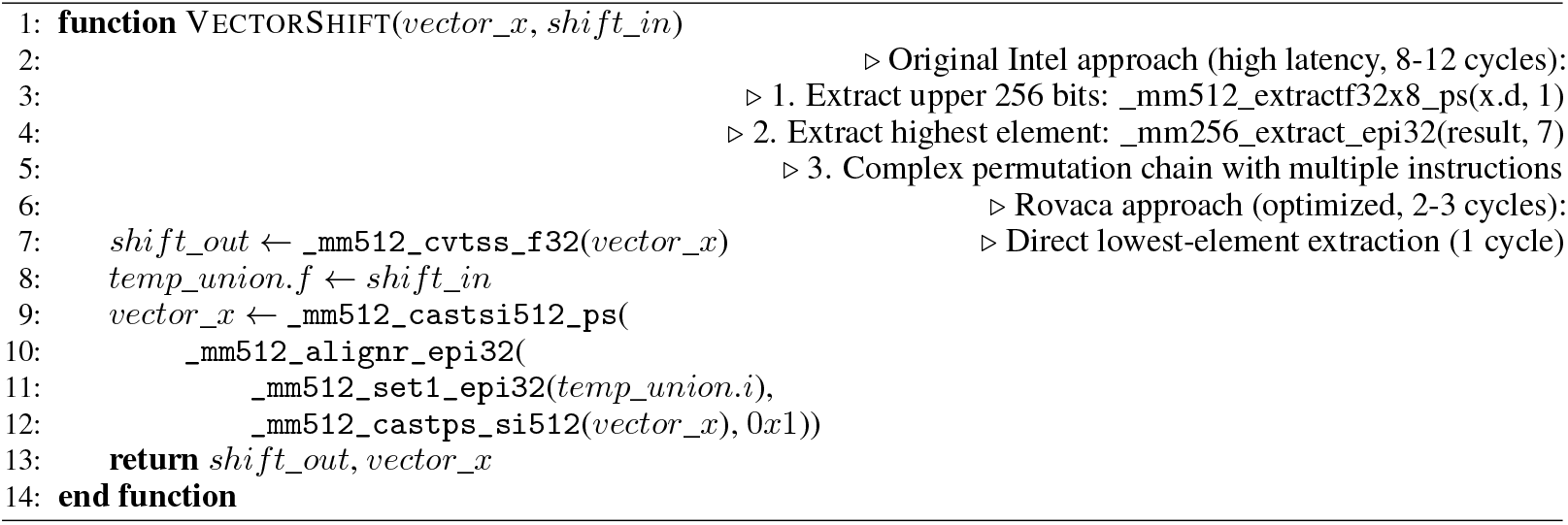

- **Pipeline Efficiency**: Elimination of cross-lane dependencies improves instruction-level parallelism
- **Cache Efficiency**: Sequential memory access patterns during loading improve spatial locality

Critically, this optimization maintains complete mathematical equivalence with the original algorithm. The data reversal is a **pure storage transformation** that:

- Preserves all numerical computations
- Maintains algorithmic dependencies
- Ensures identical mathematical results
- Does not affect convergence properties or numerical stability

This optimization demonstrates a general principle for SIMD optimization: **co-designing data layout with micro-architectural characteristics** can yield substantial performance improvements without algorithmic modification. The technique is particularly relevant for iterative algorithms requiring frequent data extraction, applications with regular data access patterns, and workloads where micro-optimizations accumulate to significant performance gains.

This innovation represents a sophisticated understanding of the interplay between algorithm design, data structure organization, and modern processor micro-architecture, achieving substantial performance improvements through minimal code changes while preserving complete algorithmic fidelity.

#### 4.4.1 Pre-computed Likelihood for Efficient Calculation

To further reduce redundant computations, Rovaca pre-calculates likelihood values. Since Base Quality (BASEQ) scores fall within a fixed range (0-59), we first count the occurrences of each BASEQ value in a read. These counts are then used to retrieve pre-computed, log-transformed likelihoods from a lookup table. This approach avoids repetitive probability calculations for each base and significantly accelerates both the active region calculation and final genotyping steps while maintaining accuracy.

### 4.5 Performance and Accuracy Evaluation

We used the hap.py toolkit to benchmark the accuracy of Rovaca’s variant calls against the GIAB truth sets. The following standard metrics were calculated for both SNPs and INDELs:

- **Precision:** The proportion of true variants among all called variants. 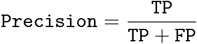
- **Recall (Sensitivity):** The proportion of true variants that were correctly identified. 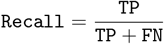
- **F1-score:** The harmonic mean of Precision and Recall, providing a single metric for accuracy. 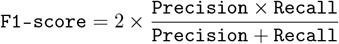

Where TP = True Positives, FP = False Positives, and FN = False Negatives.

## 5 Data Availability

Rovaca is released as open-source software and is freely available at https://github.com/ZephyRoy/Rovaca.

## 6 Supplementary Information

Supplementary information is available for this paper.

- Supplemental Table 1: Detailed runtime for different thread counts on HG001 35X WGS dataset.
- Supplemental Table 2: Detailed performance metrics for WGS and WES datasets across all samples.
- Supplemental Table 3: Detailed accuracy metrics (Precision, Recall, F1-score) for all samples and variant types for WGS.
- Supplemental Table 4: Detailed accuracy metrics (Precision, Recall, F1-score) for all samples and variant types for WES.

## References

[1] Euan A. Ashley et al. “Clinical assessment incorporating a personal genome”. In: The Lancet 375.9725 (2010), pp. 1525–1535. DOI: 10.1016/S0140-6736(10)60432-7.

[2] Adam Cornish and Chittibabu Guda. “A Comparison of Variant Calling Pipelines Using Genome in a Bottle as a Reference”. In: BioMed Research International 2015.1 (2015), p. 456479. DOI: 10.1155/2015/456479.

[3] Mark A. DePristo et al. “A framework for variation discovery and genotyping using next-generation DNA sequencing data”. In: Nature Genetics 43.5 (2011), pp. 491–498. DOI: 10.1038/ng.806.

[4] Frederick E. Dewey et al. “Clinical Interpretation and Implications of Whole-Genome Sequencing”. In: JAMA 311.10 (2014), pp. 1035–1045. DOI: 10.1001/jama.2014.1717.

[5] Genome in a Bottle. Genome in a Bottle. URL: https://www.nist.gov/programs-projects/genome-bottle (visited on 08/09/2025).

[6] Sara Goodwin, John D. McPherson, and W. Richard McCombie. “Coming of age: ten years of next-generation sequencing technologies”. In: Nature Reviews Genetics 17.6 (2016), pp. 333–351. DOI: 10.1038/nrg.2016.49.

[7] Illumina. Haplotype Comparison Tools. URL: https://github.com/Illumina/hap.py (visited on 08/09/2025).

[8] Katherine I. Kendig et al. “Sentieon DNASeq Variant Calling Workflow Demonstrates Strong Computational Performance and Accuracy”. In: Frontiers in Genetics 10 (2019), p. 736. DOI: 10.3389/fgene.2019.00736.

[9] Sangtae Kim et al. “Strelka2: fast and accurate variant calling for clinical sequencing applications”. In: bioRxiv (2017). DOI: 10.1101/192872.

[10] Hane Lee et al. “Clinical exome sequencing for genetic identification of rare Mendelian disorders”. In: JAMA 312.18 (2014), pp. 1880–1887. DOI: 10.1001/jama.2014.15671.

[11] Heng Li. Aligning sequence reads, clone sequences and assembly contigs with BWA-MEM. 2013. arXiv: 1303.3997 [q-bio.GN].

[12] Ryan Poplin et al. “A universal SNP and small-indel variant caller with deep neural networks”. In: Nature Biotechnology 36.10 (2018), pp. 983–987. DOI: 10.1038/nbt.4235.

[13] Kyle Retterer et al. “Clinical application of whole-exome sequencing across clinical indications”. In: Genetics in Medicine 18.7 (2016), pp. 696–704. DOI: 10.1038/gim.2015.148.

[14] The Broad Institute. Data pre-processing for variant discovery. URL: https://gatk.broadinstitute.org/hc/en-us/articles/360035535912-Data-pre-processing-for-variant-discovery (visited on 08/09/2025).

[15] The Broad Institute. Germline short variant discovery (SNPs + Indels). URL: https://gatk.broadinstitute.org/hc/en-us/articles/360035535932-Germline-short-variant-discovery-SNPs-Indels- (visited on 08/09/2025).

[16] UK Biobank Whole-Genome Sequencing Consortium. “Whole-genome sequencing of 490,640 UK Biobank participants”. eng. In: Nature (Aug. 2025). ISSN: 1476-4687. DOI: 10.1038/s41586-025-09272-9.

[17] Geraldine A. Van der Auwera et al. “From FastQ data to high confidence variant calls: the Genome Analysis Toolkit best practices pipeline”. In: Current Protocols in Bioinformatics 43.1 (2013), pp. 11.10.1–11.10.33. DOI: 10.1002/0471250953.bi1110s43.

[18] Yaping Yang et al. “Molecular findings among patients referred for clinical whole-exome sequencing”. In: JAMA 312.18 (2014), pp. 1870–1879. DOI: 10.1001/jama.2014.14601.

