## Supplemental Table 1 for "Rovaca: A highly optimized variant caller for rapid and robust germline DNA analysis"

Thread vs Runtime

| Thread | CPU | Memory | Runtime |
|---|---|---|---|
| 10 | 1015% | 12 | 1:27:20 |
| 20 | 2019% | 19 | 0:48:23 |
| 30 | 2999% | 25 | 0:36:51 |
| 40 | 3959% | 32 | 0:30:50 |
| 50 | 4918% | 37 | 0:26:18 |
| 60 | 5867% | 42 | 0:23:48 |
| 70 | 6805% | 49 | 0:22:16 |
| 80 | 7738% | 53 | 0:20:53 |
| 90 | 8636% | 58 | 0:19:33 |
| 100 | 9503% | 64 | 0:18:46 |
