## Supplemental Table 2 for "Rovaca: A highly optimized variant caller for rapid and robust germline DNA analysis"

### Performance Comparison

#### GRCh37

| Sample | | Rovaca V1.0.0 (52) | | | GATK-HC V4.6.0.0 | | |
|---|---|---|---|---|---|---|---|
| | | CPU | MEM (GB) | TIME (HH:MM:SS) | CPU | MEM (GB) | TIME (HH:MM:SS) |
| WGS | HG001 | 4721% | 27.92 | 0:19:29 | 194% | 7.69 | 23:58:58 |
| | HG002 | 4744% | 29.86 | 0:20:03 | 198% | 8.16 | 22:47:05 |
| | HG003 | 4800% | 29.49 | 0:19:19 | 198% | 6.4 | 22:17:09 |
| | HG004 | 4721% | 27.35 | 0:18:14 | 195% | 6.33 | 23:05:42 |
| | HG005 | 3826% | 27.49 | 0:18:07 | 205% | 6.49 | 18:49:36 |
| | HG006 | 3666% | 28.65 | 0:21:07 | 201% | 5.82 | 20:22:46 |
| | HG007 | 3651% | 27.23 | 0:22:14 | 199% | 8 | 21:05:53 |
| WES | HG001 | 2413% | 11.64 | 0:01:24 | 159% | 1.63 | 0:41:23 |
| | HG002 | 2281% | 10.53 | 0:01:19 | 159% | 1.51 | 0:37:58 |
| | HG003 | 2114% | 9.71 | 0:01:24 | 157% | 1.38 | 0:39:07 |
| | HG004 | 2389% | 11.51 | 0:01:21 | 158% | 1.52 | 0:41:18 |
| | HG005 | 2325% | 11.09 | 0:01:19 | 157% | 1.33 | 0:39:41 |
| | HG006 | 1690% | 9.01 | 0:01:07 | 143% | 1.21 | 0:35:04 |
| | HG007 | 1728% | 9.45 | 0:01:06 | 142% | 1.2 | 0:36:35 |

#### GRCh38

| Sample | | Rovaca V1.0.0 (52) | | | GATK-HC V4.6.0.0 | | |
|---|---|---|---|---|---|---|---|
| | | CPU | MEM (GB) | TIME (HH:MM:SS) | CPU | MEM (GB) | TIME (HH:MM:SS) |
| WGS | HG001 | 4364% | 26.8 | 0:17:10 | 193% | 7.31 | 22:32:55 |
| | HG002 | 4271% | 25.78 | 0:18:28 | 198% | 8.24 | 21:45:51 |
| | HG003 | 4504% | 27.86 | 0:17:01 | 197% | 7.39 | 21:36:52 |
| WES | HG001 | 2166% | 10.14 | 0:01:25 | 155% | 1.31 | 0:40:02 |
| | HG002 | 2147% | 10.68 | 0:01:23 | 155% | 7:40:48 | 0:38:45 |
| | HG003 | 2056% | 9.76 | 0:01:24 | 155% | 9:50:24 | 0:38:45 |
| | HG004 | 2163% | 10.27 | 0:01:22 | 153% | 6:57:36 | 0:40:31 |
| | HG005 | 2093% | 9.56 | 0:01:20 | 152% | 12:14:24 | 0:39:15 |
| | HG006 | 1598% | 8.73 | 0:01:09 | 143% | 3:50:24 | 0:33:06 |
| | HG007 | 1595% | 9.71 | 0:01:09 | 141% | 5:16:48 | 0:35:20 |
