## Supplemental Table 3 for "Rovaca: A highly optimized variant caller for rapid and robust germline DNA analysis"

### Accuracy Comparison for WGS

GRCh37

| HG001 | Type | TRUTH.TOTAL | TRUTH.TP | TRUTH.FN | QUERY.TOTAL | QUERY.FP | QUERY.UNK | METRIC.Recall | METRIC.Precision | METRIC.F1_Score |
|---|---|---|---|---|---|---|---|---|---|---|
| GATK-HC V4.6.0.0 | SNP | 3209315 | 3204500 | 4815 | 4009568 | 23262 | 781029 | 99.8500% | 99.2795% | 99.5639% |
| Rovaca V1.0.0 | SNP | 3209315 | 3204500 | 4815 | 4009527 | 23254 | 780996 | 99.8500% | 99.2797% | 99.5640% |
| GATK-HC V4.6.0.0 | INDEL | 481840 | 478417 | 3423 | 996726 | 3071 | 492797 | 99.2896% | 99.3906% | 99.3401% |
| Rovaca V1.0.0 | INDEL | 481840 | 478417 | 3423 | 996738 | 3072 | 492808 | 99.2896% | 99.3904% | 99.3400% |

| HG002 | Type | TRUTH.TOTAL | TRUTH.TP | TRUTH.FN | QUERY.TOTAL | QUERY.FP | QUERY.UNK | METRIC.Recall | METRIC.Precision | METRIC.F1_Score |
|---|---|---|---|---|---|---|---|---|---|---|
| GATK-HC V4.6.0.0 | SNP | 3352679 | 3328324 | 24355 | 3954318 | 55119 | 569894 | 99.2736% | 98.3714% | 98.8204% |
| Rovaca V1.0.0 | SNP | 3352679 | 3328323 | 24356 | 3954240 | 55113 | 569823 | 99.2735% | 98.3716% | 98.8205% |
| GATK-HC V4.6.0.0 | INDEL | 522389 | 519042 | 3347 | 986689 | 5489 | 439913 | 99.3593% | 98.9961% | 99.1774% |
| Rovaca V1.0.0 | INDEL | 522389 | 519042 | 3347 | 986717 | 5488 | 439942 | 99.3593% | 98.9963% | 99.1775% |

| HG003 | Type | TRUTH.TOTAL | TRUTH.TP | TRUTH.FN | QUERY.TOTAL | QUERY.FP | QUERY.UNK | METRIC.Recall | METRIC.Precision | METRIC.F1_Score |
|---|---|---|---|---|---|---|---|---|---|---|
| GATK-HC V4.6.0.0 | SNP | 3314143 | 3289886 | 24257 | 3922218 | 54402 | 577076 | 99.2681% | 98.3737% | 98.8189% |
| Rovaca V1.0.0 | SNP | 3314143 | 3289887 | 24256 | 3922291 | 54410 | 577140 | 99.2681% | 98.3735% | 98.8188% |
| GATK-HC V4.6.0.0 | INDEL | 500793 | 497419 | 3374 | 977504 | 5390 | 453555 | 99.3263% | 98.9713% | 99.1485% |
| Rovaca V1.0.0 | INDEL | 500793 | 497419 | 3374 | 977527 | 5394 | 453574 | 99.3263% | 98.9705% | 99.1481% |

| HG004 | Type | TRUTH.TOTAL | TRUTH.TP | TRUTH.FN | QUERY.TOTAL | QUERY.FP | QUERY.UNK | METRIC.Recall | METRIC.Precision | METRIC.F1_Score |
|---|---|---|---|---|---|---|---|---|---|---|
| GATK-HC V4.6.0.0 | SNP | 3340471 | 3313157 | 27314 | 3950191 | 55979 | 580196 | 99.1823% | 98.3389% | 98.7588% |
| Rovaca V1.0.0 | SNP | 3340471 | 3313157 | 27314 | 3950281 | 55978 | 580287 | 99.1823% | 98.3389% | 98.7588% |
| GATK-HC V4.6.0.0 | INDEL | 508274 | 503873 | 4401 | 971958 | 5525 | 440567 | 99.1341% | 98.9603% | 99.0471% |
| Rovaca V1.0.0 | INDEL | 508274 | 503873 | 4401 | 971995 | 5526 | 440603 | 99.1341% | 98.9601% | 99.0470% |

| HG005 | Type | TRUTH.TOTAL | TRUTH.TP | TRUTH.FN | QUERY.TOTAL | QUERY.FP | QUERY.UNK | METRIC.Recall | METRIC.Precision | METRIC.F1_Score |
|---|---|---|---|---|---|---|---|---|---|---|
| GATK-HC V4.6.0.0 | SNP | 3265971 | 3227068 | 38903 | 3868064 | 50548 | 589863 | 98.8088% | 98.4581% | 98.6331% |
| Rovaca V1.0.0 | SNP | 3265971 | 3227068 | 38903 | 3868026 | 50554 | 589819 | 98.8088% | 98.4579% | 98.6330% |
| GATK-HC V4.6.0.0 | INDEL | 413821 | 409500 | 4321 | 958210 | 5308 | 530062 | 98.9558% | 98.7602% | 98.8579% |
| Rovaca V1.0.0 | INDEL | 413821 | 409500 | 4321 | 958219 | 5309 | 530070 | 98.9558% | 98.7600% | 98.8578% |

| HG006 | Type | TRUTH.TOTAL | TRUTH.TP | TRUTH.FN | QUERY.TOTAL | QUERY.FP | QUERY.UNK | METRIC.Recall | METRIC.Precision | METRIC.F1_Score |
|---|---|---|---|---|---|---|---|---|---|---|
| GATK-HC V4.6.0.0 | SNP | 3275380 | 3238206 | 37174 | 3851231 | 50014 | 562276 | 98.8650% | 98.4793% | 98.6718% |
| Rovaca V1.0.0 | SNP | 3275380 | 3238207 | 37173 | 3851340 | 50018 | 562380 | 98.8651% | 98.4792% | 98.6718% |
| GATK-HC V4.6.0.0 | INDEL | 446174 | 441552 | 4622 | 953434 | 5224 | 491436 | 98.9641% | 98.8693% | 98.9166% |
| Rovaca V1.0.0 | INDEL | 446174 | 441552 | 4622 | 953433 | 5222 | 491437 | 98.9641% | 98.8697% | 98.9169% |

| HG007 | Type | TRUTH.TOTAL | TRUTH.TP | TRUTH.FN | QUERY.TOTAL | QUERY.FP | QUERY.UNK | METRIC.Recall | METRIC.Precision | METRIC.F1_Score |
|---|---|---|---|---|---|---|---|---|---|---|

Accuracy Comparison for WGS

| | | | | | | | | | | |
|---|---|---|---|---|---|---|---|---|---|---|
| **GATK-HC V4.6.0.0** | SNP | 3271790 | 3238492 | 33298 | 3880799 | 41218 | 600422 | 98.9823% | 98.7435% | 98.8627% |
| **Rovaca V1.0.0** | SNP | 3271790 | 3238493 | 33297 | 3880790 | 41223 | 600407 | 98.9823% | 98.7433% | 98.8627% |
| **GATK-HC V4.6.0.0** | INDEL | 439073 | 434911 | 4162 | 957654 | 4375 | 503057 | 99.0521% | 99.0376% | 99.0449% |
| **Rovaca V1.0.0** | INDEL | 439073 | 434911 | 4162 | 957664 | 4375 | 503067 | 99.0521% | 99.0376% | 99.0449% |

**GRCh38**

| HG001 | Type | TRUTH.TOTAL | TRUTH.TP | TRUTH.FN | QUERY.TOTAL | QUERY.FP | QUERY.UNK | METRIC.Recall | METRIC.Precision | METRIC.F1_Score |
|---|---|---|---|---|---|---|---|---|---|---|
| **GATK-HC V4.6.0.0** | SNP | 3254352 | 3230561 | 23791 | 3864690 | 27381 | 605988 | 99.2689% | 99.1598% | 99.2143% |
| **Rovaca V1.0.0** | SNP | 3254352 | 3230561 | 23791 | 3864708 | 27382 | 606005 | 99.2689% | 99.1597% | 99.2143% |
| **GATK-HC V4.6.0.0** | INDEL | 467684 | 463949 | 3735 | 953591 | 2786 | 468575 | 99.2014% | 99.4256% | 99.3134% |
| **Rovaca V1.0.0** | INDEL | 467684 | 463949 | 3735 | 953594 | 2786 | 468578 | 99.2014% | 99.4256% | 99.3134% |

| HG002 | Type | TRUTH.TOTAL | TRUTH.TP | TRUTH.FN | QUERY.TOTAL | QUERY.FP | QUERY.UNK | METRIC.Recall | METRIC.Precision | METRIC.F1_Score |
|---|---|---|---|---|---|---|---|---|---|---|
| **GATK-HC V4.6.0.0** | SNP | 3365115 | 3338895 | 26220 | 3925133 | 26265 | 558956 | 99.2208% | 99.2197% | 99.2203% |
| **Rovaca V1.0.0** | SNP | 3365115 | 3338894 | 26221 | 3925131 | 26267 | 558953 | 99.2208% | 99.2197% | 99.2202% |
| **GATK-HC V4.6.0.0** | INDEL | 525466 | 521993 | 3473 | 979829 | 2650 | 432726 | 99.3391% | 99.5156% | 99.4273% |
| **Rovaca V1.0.0** | INDEL | 525466 | 521993 | 3473 | 979839 | 2650 | 432736 | 99.3391% | 99.5156% | 99.4273% |

| HG003 | Type | TRUTH.TOTAL | TRUTH.TP | TRUTH.FN | QUERY.TOTAL | QUERY.FP | QUERY.UNK | METRIC.Recall | METRIC.Precision | METRIC.F1_Score |
|---|---|---|---|---|---|---|---|---|---|---|
| **GATK-HC V4.6.0.0** | SNP | 3327481 | 3300741 | 26740 | 3896176 | 26681 | 567845 | 99.1964% | 99.1984% | 99.1974% |
| **Rovaca V1.0.0** | SNP | 3327481 | 3300742 | 26739 | 3896158 | 26681 | 567826 | 99.1964% | 99.1984% | 99.1974% |
| **GATK-HC V4.6.0.0** | INDEL | 504497 | 501051 | 3446 | 971221 | 2719 | 446113 | 99.3169% | 99.4822% | 99.3995% |
| **Rovaca V1.0.0** | INDEL | 504497 | 501051 | 3446 | 971230 | 2719 | 446122 | 99.3169% | 99.4822% | 99.3995% |
