## Supplemental Table 4 for "Rovaca: A highly optimized variant caller for rapid and robust germline DNA analysis"

Accuracy Comparison for WES

GRCh37

| HG001 | Type | TRUTH.TOTAL | TRUTH.TP | TRUTH.FN | QUERY.TOTAL | QUERY.FP | QUERY.UNK | METRIC.Recall | METRIC.Precision | METRIC.F1_Score |
|---|---|---|---|---|---|---|---|---|---|---|
| GATK-HC V4.6.0.0 | SNP | 20497 | 19595 | 902 | 24913 | 283 | 5037 | 95.5994% | 98.5762% | 97.0649% |
| Rovaca V1.0.0 | SNP | 20497 | 19595 | 902 | 24917 | 287 | 5037 | 95.5994% | 98.5563% | 97.0553% |
| GATK-HC V4.6.0.0 | INDEL | 626 | 534 | 92 | 1123 | 144 | 432 | 85.3035% | 79.1606% | 82.1174% |
| Rovaca V1.0.0 | INDEL | 626 | 534 | 92 | 1124 | 145 | 432 | 85.3035% | 79.0462% | 82.0558% |

| HG002 | Type | TRUTH.TOTAL | TRUTH.TP | TRUTH.FN | QUERY.TOTAL | QUERY.FP | QUERY.UNK | METRIC.Recall | METRIC.Precision | METRIC.F1_Score |
|---|---|---|---|---|---|---|---|---|---|---|
| GATK-HC V4.6.0.0 | SNP | 22743 | 21128 | 1615 | 24588 | 696 | 2764 | 92.8989% | 96.8109% | 94.8145% |
| Rovaca V1.0.0 | SNP | 22743 | 21128 | 1615 | 24587 | 696 | 2763 | 92.8989% | 96.8109% | 94.8145% |
| GATK-HC V4.6.0.0 | INDEL | 682 | 580 | 102 | 1091 | 156 | 350 | 85.0440% | 78.9474% | 81.8824% |
| Rovaca V1.0.0 | INDEL | 682 | 580 | 102 | 1091 | 156 | 350 | 85.0440% | 78.9474% | 81.8824% |

| HG003 | Type | TRUTH.TOTAL | TRUTH.TP | TRUTH.FN | QUERY.TOTAL | QUERY.FP | QUERY.UNK | METRIC.Recall | METRIC.Precision | METRIC.F1_Score |
|---|---|---|---|---|---|---|---|---|---|---|
| GATK-HC V4.6.0.0 | SNP | 22589 | 21060 | 1529 | 24641 | 580 | 3000 | 93.2312% | 97.3199% | 95.2317% |
| Rovaca V1.0.0 | SNP | 22589 | 21060 | 1529 | 24637 | 579 | 2997 | 93.2312% | 97.3244% | 95.2338% |
| GATK-HC V4.6.0.0 | INDEL | 690 | 593 | 97 | 1110 | 132 | 377 | 85.9420% | 81.9918% | 83.9205% |
| Rovaca V1.0.0 | INDEL | 690 | 593 | 97 | 1107 | 132 | 374 | 85.9420% | 81.9918% | 83.9205% |

| HG004 | Type | TRUTH.TOTAL | TRUTH.TP | TRUTH.FN | QUERY.TOTAL | QUERY.FP | QUERY.UNK | METRIC.Recall | METRIC.Precision | METRIC.F1_Score |
|---|---|---|---|---|---|---|---|---|---|---|
| GATK-HC V4.6.0.0 | SNP | 22543 | 20976 | 1567 | 24779 | 631 | 3173 | 93.0488% | 97.0795% | 95.0215% |
| Rovaca V1.0.0 | SNP | 22543 | 20976 | 1567 | 24784 | 636 | 3173 | 93.0488% | 97.0571% | 95.0107% |
| GATK-HC V4.6.0.0 | INDEL | 643 | 557 | 86 | 1094 | 157 | 367 | 86.6252% | 78.4044% | 82.3100% |
| Rovaca V1.0.0 | INDEL | 643 | 557 | 86 | 1095 | 157 | 368 | 86.6252% | 78.4044% | 82.3100% |

| HG005 | Type | TRUTH.TOTAL | TRUTH.TP | TRUTH.FN | QUERY.TOTAL | QUERY.FP | QUERY.UNK | METRIC.Recall | METRIC.Precision | METRIC.F1_Score |
|---|---|---|---|---|---|---|---|---|---|---|
| GATK-HC V4.6.0.0 | SNP | 22543 | 20975 | 1568 | 24829 | 628 | 3226 | 93.0444% | 97.0930% | 95.0256% |
| Rovaca V1.0.0 | SNP | 22543 | 20975 | 1568 | 24829 | 626 | 3228 | 93.0444% | 97.1020% | 95.0299% |
| GATK-HC V4.6.0.0 | INDEL | 615 | 529 | 86 | 1113 | 123 | 454 | 86.0163% | 81.3354% | 83.6103% |
| Rovaca V1.0.0 | INDEL | 615 | 529 | 86 | 1127 | 136 | 455 | 86.0163% | 79.7619% | 82.7711% |

| HG006 | Type | TRUTH.TOTAL | TRUTH.TP | TRUTH.FN | QUERY.TOTAL | QUERY.FP | QUERY.UNK | METRIC.Recall | METRIC.Precision | METRIC.F1_Score |
|---|---|---|---|---|---|---|---|---|---|---|
| GATK-HC V4.6.0.0 | SNP | 22306 | 20644 | 1662 | 23933 | 615 | 2673 | 92.5491% | 97.1072% | 94.7734% |
| Rovaca V1.0.0 | SNP | 22306 | 20644 | 1662 | 23925 | 613 | 2667 | 92.5491% | 97.1164% | 94.7777% |
| GATK-HC V4.6.0.0 | INDEL | 615 | 522 | 93 | 964 | 60 | 373 | 84.8780% | 89.8477% | 87.2922% |
| Rovaca V1.0.0 | INDEL | 615 | 522 | 93 | 964 | 60 | 373 | 84.8780% | 89.8477% | 87.2922% |

| HG007 | Type | TRUTH.TOTAL | TRUTH.TP | TRUTH.FN | QUERY.TOTAL | QUERY.FP | QUERY.UNK | METRIC.Recall | METRIC.Precision | METRIC.F1_Score |
|---|---|---|---|---|---|---|---|---|---|---|

### Accuracy Comparison for WES

| | | | | | | | | | | |
|---|---|---|---|---|---|---|---|---|---|---|
| GATK-HC V4.6.0.0 | SNP | 22322 | 20604 | 1718 | 24206 | 478 | 3123 | 92.3036% | 97.7328% | 94.9406% |
| Rovaca V1.0.0 | SNP | 22322 | 20604 | 1718 | 24204 | 476 | 3123 | 92.3036% | 97.7420% | 94.9450% |
| GATK-HC V4.6.0.0 | INDEL | 589 | 499 | 90 | 925 | 56 | 366 | 84.7199% | 89.9821% | 87.2717% |
| Rovaca V1.0.0 | INDEL | 589 | 499 | 90 | 937 | 68 | 366 | 84.7199% | 88.0911% | 86.3726% |

**GRCh38**

| HG001 | Type | TRUTH.TOTAL | TRUTH.TP | TRUTH.FN | QUERY.TOTAL | QUERY.FP | QUERY.UNK | METRIC.Recall | METRIC.Precision | METRIC.F1_Score |
|---|---|---|---|---|---|---|---|---|---|---|
| GATK-HC V4.6.0.0 | SNP | 21973 | 20511 | 1462 | 24149 | 485 | 3153 | 93.3464% | 97.6900% | 95.4688% |
| Rovaca V1.0.0 | SNP | 21973 | 20511 | 1462 | 24149 | 485 | 3153 | 93.3464% | 97.6900% | 95.4688% |
| GATK-HC V4.6.0.0 | INDEL | 549 | 478 | 71 | 959 | 135 | 337 | 87.0674% | 78.2958% | 82.4490% |
| Rovaca V1.0.0 | INDEL | 549 | 478 | 71 | 959 | 135 | 337 | 87.0674% | 78.2958% | 82.4490% |

| HG002 | Type | TRUTH.TOTAL | TRUTH.TP | TRUTH.FN | QUERY.TOTAL | QUERY.FP | QUERY.UNK | METRIC.Recall | METRIC.Precision | METRIC.F1_Score |
|---|---|---|---|---|---|---|---|---|---|---|
| GATK-HC V4.6.0.0 | SNP | 22708 | 21082 | 1626 | 24044 | 515 | 2450 | 92.8395% | 97.6151% | 95.1674% |
| Rovaca V1.0.0 | SNP | 22708 | 21082 | 1626 | 24044 | 515 | 2450 | 92.8395% | 97.6151% | 95.1674% |
| GATK-HC V4.6.0.0 | INDEL | 561 | 483 | 78 | 934 | 138 | 306 | 86.0963% | 78.0255% | 81.8624% |
| Rovaca V1.0.0 | INDEL | 561 | 483 | 78 | 934 | 138 | 306 | 86.0963% | 78.0255% | 81.8624% |

| HG003 | Type | TRUTH.TOTAL | TRUTH.TP | TRUTH.FN | QUERY.TOTAL | QUERY.FP | QUERY.UNK | METRIC.Recall | METRIC.Precision | METRIC.F1_Score |
|---|---|---|---|---|---|---|---|---|---|---|
| GATK-HC V4.6.0.0 | SNP | 22576 | 21072 | 1504 | 24114 | 436 | 2604 | 93.3381% | 97.9730% | 95.5994% |
| Rovaca V1.0.0 | SNP | 22576 | 21072 | 1504 | 24113 | 436 | 2603 | 93.3381% | 97.9730% | 95.5994% |
| GATK-HC V4.6.0.0 | INDEL | 570 | 499 | 71 | 945 | 123 | 315 | 87.5439% | 80.4762% | 83.8614% |
| Rovaca V1.0.0 | INDEL | 570 | 499 | 71 | 945 | 123 | 315 | 87.5439% | 80.4762% | 83.8614% |

| HG004 | Type | TRUTH.TOTAL | TRUTH.TP | TRUTH.FN | QUERY.TOTAL | QUERY.FP | QUERY.UNK | METRIC.Recall | METRIC.Precision | METRIC.F1_Score |
|---|---|---|---|---|---|---|---|---|---|---|
| GATK-HC V4.6.0.0 | SNP | 22475 | 20875 | 1600 | 24167 | 437 | 2856 | 92.8810% | 97.9494% | 95.3479% |
| Rovaca V1.0.0 | SNP | 22475 | 20875 | 1600 | 24166 | 436 | 2856 | 92.8810% | 97.9540% | 95.3501% |
| GATK-HC V4.6.0.0 | INDEL | 528 | 470 | 58 | 936 | 130 | 322 | 89.0152% | 78.8274% | 83.6121% |
| Rovaca V1.0.0 | INDEL | 528 | 470 | 58 | 935 | 130 | 321 | 89.0152% | 78.8274% | 83.6121% |

| HG005 | Type | TRUTH.TOTAL | TRUTH.TP | TRUTH.FN | QUERY.TOTAL | QUERY.FP | QUERY.UNK | METRIC.Recall | METRIC.Precision | METRIC.F1_Score |
|---|---|---|---|---|---|---|---|---|---|---|
| GATK-HC V4.6.0.0 | SNP | 22507 | 21010 | 1497 | 24259 | 514 | 2735 | 93.3487% | 97.6120% | 95.4328% |
| Rovaca V1.0.0 | SNP | 22507 | 21010 | 1497 | 24257 | 515 | 2732 | 93.3487% | 97.6074% | 95.4306% |
| GATK-HC V4.6.0.0 | INDEL | 508 | 449 | 59 | 948 | 112 | 382 | 88.3858% | 80.2120% | 84.1008% |
| Rovaca V1.0.0 | INDEL | 508 | 449 | 59 | 949 | 113 | 382 | 88.3858% | 80.0705% | 84.0230% |

| HG006 | Type | TRUTH.TOTAL | TRUTH.TP | TRUTH.FN | QUERY.TOTAL | QUERY.FP | QUERY.UNK | METRIC.Recall | METRIC.Precision | METRIC.F1_Score |
|---|---|---|---|---|---|---|---|---|---|---|
| GATK-HC V4.6.0.0 | SNP | 22268 | 20585 | 1683 | 23414 | 445 | 2383 | 92.4421% | 97.8841% | 95.0853% |

Accuracy Comparison for WES

| | | | | | | | | | | |
|---|---|---|---|---|---|---|---|---|---|---|
| **Rovaca V1.0.0** | SNP | 22268 | 20585 | 1683 | 23417 | 445 | 2386 | 92.4421% | 97.8841% | 95.0853% |
| **GATK-HC V4.6.0.0** | INDEL | 492 | 427 | 65 | 839 | 46 | 357 | 86.7886% | 90.4564% | 88.5846% |
| **Rovaca V1.0.0** | INDEL | 492 | 427 | 65 | 837 | 46 | 355 | 86.7886% | 90.4564% | 88.5846% |

| HG007 | Type | TRUTH.TOTAL | TRUTH.TP | TRUTH.FN | QUERY.TOTAL | QUERY.FP | QUERY.UNK | METRIC.Recall | METRIC.Precision | METRIC.F1_Score |
|---|---|---|---|---|---|---|---|---|---|---|
| **GATK-HC V4.6.0.0** | SNP | 22416 | 20681 | 1735 | 23694 | 417 | 2595 | 92.2600% | 98.0236% | 95.0545% |
| **Rovaca V1.0.0** | SNP | 22416 | 20681 | 1735 | 23695 | 417 | 2596 | 92.2600% | 98.0236% | 95.0545% |
| **GATK-HC V4.6.0.0** | INDEL | 478 | 411 | 67 | 778 | 49 | 314 | 85.9833% | 89.4397% | 87.6774% |
| **Rovaca V1.0.0** | INDEL | 478 | 411 | 67 | 778 | 49 | 314 | 85.9833% | 89.4397% | 87.6774% |
